# Accurate bulk quantitation of droplet digital PCR

**DOI:** 10.1101/2021.01.13.424628

**Authors:** Chen Sun, Leqian Liu, Harish N. Vasudevan, Kai-Chun Chang, Adam R. Abate

## Abstract

Droplet digital PCR provides superior accuracy in nucleic acid quantitation. The requirement of microfluidics to generate and analyze the emulsions, however, is a barrier to its adoption, particularly in low resource or clinical settings. Here, we report a novel method to prepare ddPCR droplets by vortexing and readout the results by bulk analysis of recovered amplicons. We demonstrate the approach by accurately quantitating SARS-CoV-2 sequences using entirely bulk processing and no microfluidics. Our approach for quantitating reactions should extend to all digital assays that generate amplicons, including digital PCR and LAMP conducted in droplets, microchambers, or nanoliter wells. More broadly, our approach combines important attributes of ddPCR, including enhanced accuracy and robustness to inhibition, with the high-volume sample processing ability of quantitative PCR.

## Introduction

The quantitation of nucleic acids is important for basic science and clinical applications. Quantitative PCR (qPCR) measures target concentration by monitoring the exponential rise of amplicons and is the gold standard due to its specificity and superb sensitivity^1^. By contrast, digital PCR (dPCR) subdivides the sample such that partitions contain one or no target molecule; after end-point amplification, positives are enumerated, yielding the target concentration^2–4^. Digital PCR affords numerous advantages over qPCR, including absolute quantitation and enhanced accuracy for small concentration changes, making it especially valuable for clinical applications^5–7^. It has secondary benefits, including increased resistance to inhibition^8–9^ and the ability to differentiate intact from fragmented molecules^10–11^, which are valuable in the identification of viable pathogens in minimally processed samples^12–13^.

Droplet digital PCR (ddPCR) uses microfluidics to partition samples in water droplets suspended in oil. While the approach is superbly accurate, the requirement of microfluidics is a barrier to its adoption, making it costly compared to qPCR, and difficult to integrate into clinical labs using standardized well plate formats. Particle-templated emulsification (PTE) partitions samples without microfluidics; the resultant emulsions are similar in monodispersity to microfluidically-generated ones and, thus, can be used to conduct most droplet assays, including ddPCR^14^. While removal of microfluidic droplet generation is a great simplification, subsequent quantification still requires a custom droplet reader, negating much of the advantage^15–17^. To realize the benefits of ddPCR in settings in which microfluidic instrumentation is impractical, a new approach for enumerating positive droplets that uses only common laboratory equipment and methods is needed.

In this paper, we demonstrate accurate bulk quantitation of droplet digital PCR with common lab equipment. To partition the samples, we use bulk homogenization with a vortexer. To quantitate the samples, we compare different methods for bulk enumeration of positive droplets, including fluorescence, gel electrophoresis, and qPCR. Of these, we find Qpcr detection of droplet products yields the highest sensitivity and accuracy over the widest dynamic range. Thus, our approach combines important attributes of ddPCR, including enhanced accuracy and robustness to inhibition, with the accessibility and scalability of bulk processing in well plates. We demonstrate the method by using it to quantify SARS-CoV-2 nucleic acids. While we focus on droplet dPCR, the principles of our bulk quantitation should apply to any dPCR approach in which amplicons can be recovered from the partitions and analyzed, including nanoliter well and microchamber technologies^3, 18–20^.

## Results and Discussion

An important advantage of ddPCR over qPCR is its ability to accurately quantify small differences in target concentration, especially near the detection limit of the assay^15, 21^. This benefit arises from the linear nature of ddPCR. Because qPCR is exponential, stochasticity in reaction initiation amplifies over cycles to limit the precision with which small differences in target concentration can be measured. By contrast, when cycled to endpoint, irrespective of when each droplet amplification initiates, the number of positive droplets in ddPCR is directly proportional to the number of input target molecules (Figure 1a). This allows accurate measurement of target concentration,

**Fig 1.**
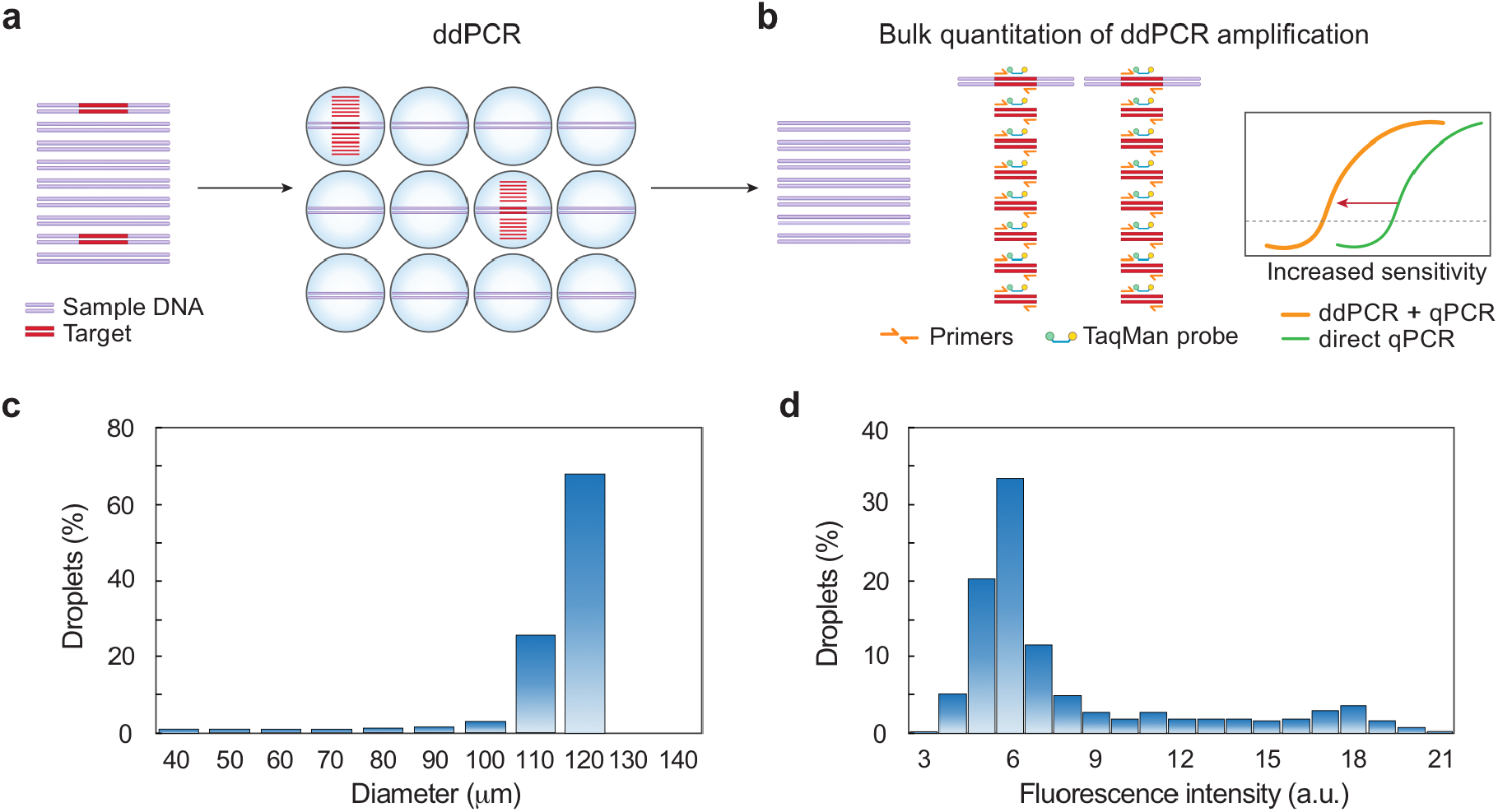
Schematic of workflow for bulk quantitation of ddPCR. (**a**) A mixed DNA sample is emulsified and processed for ddPCR. The target molecules are amplified in individual droplets. (**b**) The number of target molecules in the starting sample is proportional to the amount of amplification products, which are quantified by bulk measurement. qPCR quantification of the ddPCR products shows enhanced sensitivity compared to direct qPCR by elevating the qPCR signal. (**c**) Size distribution of microfluidic emulsions (*n* = 950) shows microfluidics generates monodispersed emulsions. (**d**) Fluorescence intensity distribution of microfluidic emulsions (*n* = 540) indicates that the assay has nonzero background.

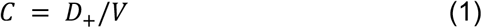

where *C* is the target concentration, *D*_+_ the number of positive droplets, and V the total volume of the sample, and is the basis of ddPCR’s ability to obtain an “absolute” count of target molecules, while qPCR returns only relative values unless a standard curve is provided^9^. Thus, enumerating positive droplets is an essential step in ddPCR, and is typically accomplished using a droplet reader comprising a microfluidic optical instrument^16–17^. In addition to being costly, these instruments are difficult to integrate into high-volume testing because each sample must be manually processed; consequently, they are reserved primarily to settings that can bear the high labor and equipment costs^3^. A superior strategy would be to infer positive droplet number from a bulk measurement compatible with plate-processing of samples; this would significantly lower the barrier to adoption and enable high throughput processing in plates.

In principle, the total fluorescence of an emulsion provides a straightforward way to infer the number of positive droplets because it is the sum of the contributions of the positive F_+_ and negative F_−_ droplet fluorescence,

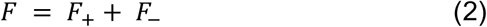

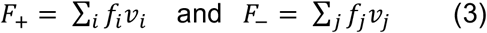

with *f*_*i*_ the fluorescence density and *v*_*i*_ the volume of the *i*th positive droplet; and *f*_*j*_ the fluorescence density and *v*_*j*_ the volume of the *j*th negative droplet. In the limit F_+_ ≫ F_.−_, and assuming each positive droplet contributes an average quantum of fluorescence 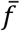 (Figure 1c), the number of positive droplets

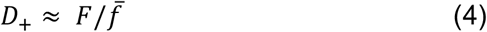

Thus, for a suitable background fluorescence, it is possible to infer D_+_ from bulk measurement of the fluorescence emerging from an emulsion^22^. Nevertheless, bulk fluorescence is a poor observable due to the optical properties of ddPCR emulsions. Unless the carrier oil is index matched to the droplets, emulsions are opaque^23–24^; the amount of signal detected from a droplet deep within the emulsion may thus differ from one near the surface. In addition, common methods for measuring fluorescence in wells read from the bottom which limits reproducibility, since collection efficiency will depend on where the emulsion is in the tube and how long it has settled before being read. Most importantly, ddPCR assays have nonzero background (*f*_*j*_ is not negligible compared to *f* _*i*_) (Figure 1d) such that the condition *F*_+_ ≫ *F*_.−_ is usually only met when the number of positive drops is large; this limits sensitivity for the most important low concentrations.

In addition to fluorescence, ddPCRs produce amplicons (Figure 1b). In principle, if similar conditions are met of low background and uniform generation from droplets, bulk measurement of amplicons should allow inference of positives in analogy to Eq. (4),

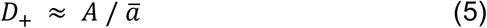

where *A* is the total number of amplicons generated by a ddPCR and ā the average number generated per positive droplet. Like total fluorescence, this approximation is justified when the number of amplicons generated by the positive droplets is much greater than by the negatives (A_+_ ≫ A_.−_). In this respect, amplicon detection is superior to fluorescence because well-designed PCRs generate few off-target products. In addition, opacity of the emulsion is not a factor, and amplicons can be measured using a variety of common and sensitive techniques, including staining, on-chip electrophoresis, and qPCR^1, 25–26^.

To investigate whether amplicon quantitation provides a suitable means for estimating D_+_ in bulk, we compare the efficacy of these methods for a dilution series of SARS-CoV-2 nucleic acids (Figure 2). We generate all ddPCR assays with commercially available microfluidics^27^, and each sample is divided into 20,000 droplets. As expected due to the high background, bulk fluorescence poorly quantifies ddPCR results, yielding a detection sensitivity of ∼320 molecules (Figure 2a). Recovering and staining DNA from the droplets and quantitating with a fluorescence reader yields a sensitivity of ∼80 molecules; this technique, however, is nonspecific and detects all recovered DNA irrespective of sequence, yielding a suboptimal background (Figure 2b). To reduce background, we target the amplicons for detection using on-chip gel electrophoresis; this allows quantitation of the peak representing the correct molecular length (Supp Fig S1). The result is an improved detection sensitivity of ∼20 molecules (Figure 2c), which is nearly as good as direct qPCR analysis of the sample, also having a sensitivity of ∼20 molecules (Figure 2d, upper points). The measurement becomes less accurate at high concentrations due to multiple targets being encapsulated in the droplets. Importantly, since electrophoresis measures the lengths of all amplicons in the sample, it is readily multiplexed by designing amplicons of different length^26^. Moreover, when performed in an emulsion, multiplexed reactions tend to be robust because products do not compete for amplification^28^.

**Fig 2.**
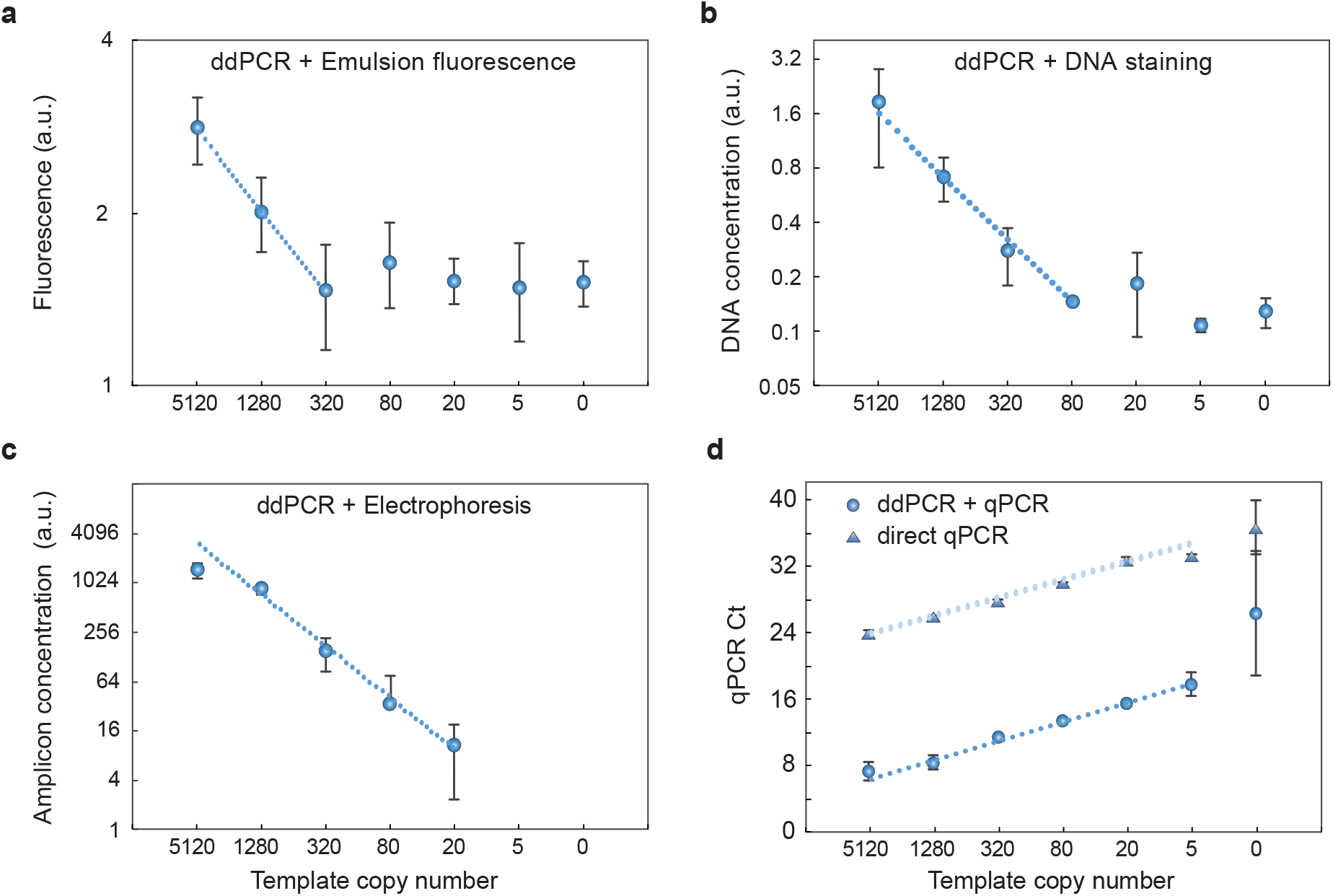
Bulk quantitation of ddPCR products. (**a**) Total fluorescence of ddPCR emulsions measured with a plate reader (Tecan); (**b**) Detection of stained total DNA recovered from ddPCR emulsions (Qubit); (**c**) Quantitation of amplicon peak with gel electrophoresis (Bioanalyzer) of ddPCR emulsions. (**d**) qPCR quantitation of ddPCR amplicons. ddPCR+qPCR shifts the qPCR *C*_*t*_ to lower cycles, allowing enhanced sensitivity compared to qPCR alone. *n* = 3, error bars represent standard deviation.

Below this detection limit, gel electrophoresis is ineffective because the recovered molecules are too few to be detected. To increase detection sensitivity further, we thus need a more sensitive amplicon quantitation approach. qPCR is a sensitive technique for quantifying nucleic acids and has the benefits of being specific and multiplexable since primers can be targeted to different sequences. As such, with qPCR of ddPCR products, we achieve a detection of just ∼5 molecules (Figure 2d). Below this, detection becomes unreliable because there are so few molecules there is large variability due to statistical loading of targets in the sample^29^. In concordance with this, we observe increased standard deviation when the sample has ∼5 targets. We observe amplification in no template controls in both direct qPCR and ddPCR+qPCR, likely due to airborne contamination or non-specific amplification. When targets are abundant, qPCR affords excellent quantitation (Figure 2d). However, direct qPCR has higher *C*_*t*_ values because it detects targets directly, while ddPCR+qPCR detects the amplicons generated by the droplets; the result is that much more DNA is present at the beginning of the qPCR analysis, yielding smaller *C*_*t*_ values. This demonstrates that bulk quantitation of ddPCR-generated amplicons, like direct droplet enumeration, is ultimately limited by statistical loading of targets in the sample and not by the assay sensitivity or accuracy.

While ddPCR+qPCR affords the best sensitivity of all methods we test and even surpasses qPCR, the requirement of microfluidics to generate the emulsions is a major limitation. Indeed, emulsions can be generated by simpler methods, including bead beating, sonication, and pipetting^14, 30–31^. Vortexing also produces emulsions, with the benefits of being simple, fast, and amenable to parallel processing. However, these bulk methods generate polydispersed emulsions in which droplet size varies substantially compared to microfluidics. While accurate ddPCR has been demonstrated in polydispersed emulsions when droplets are imaged and counted^32^, it is unclear whether this holds for bulk detection because, when cycled to endpoint, the number of amplicons generated in a droplet scales with its volume. Thus, the total number of amplicons in the recovered pool will depend on the volumes of the positive droplets, which will vary,

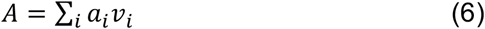

with *a*_*i*_ the amplicon concentration and *v*_*i*_ the volume produced by the *i*th positive droplet. In the limit of large *A*_+_, however, the sum can be approximated in terms of the average ā, simplifying the expression to

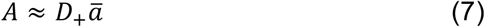

such that Eq. (5) still holds. Below this limit, statistical variation in droplet volume dominates the measurement. Where this approximation holds will thus depend on the size distribution of the droplets, such that more polydispersed emulsions will lose their quantitativeness at higher A_+_. To investigate this concept, we perform another experiment in which we quantify polydispersed ddPCR emulsions generated by vortexing (Figure 3a). As expected, the emulsions are polydispersed, though positive droplets are clearly visible (Figure 3b); in addition, the size distribution is much broader than for the microfluidic emulsion (Figure 3c). When we measure the recovered amplicons, we find excellent quantitation, with minimal error down to 20 molecules. Below this, statistical variation in droplet size increases error (Figure 3d, right) though the measurement remains quantitative down to ∼5 molecules, and nearly as good as monodispersed emulsions (Figure 2d). Furthermore, vortex-generated emulsions have smaller average droplet sizes than the microfluidic ones and, thus, the sample is subdivided into more partitions, increasing dynamic range at higher concentrations.

**Fig 3.**
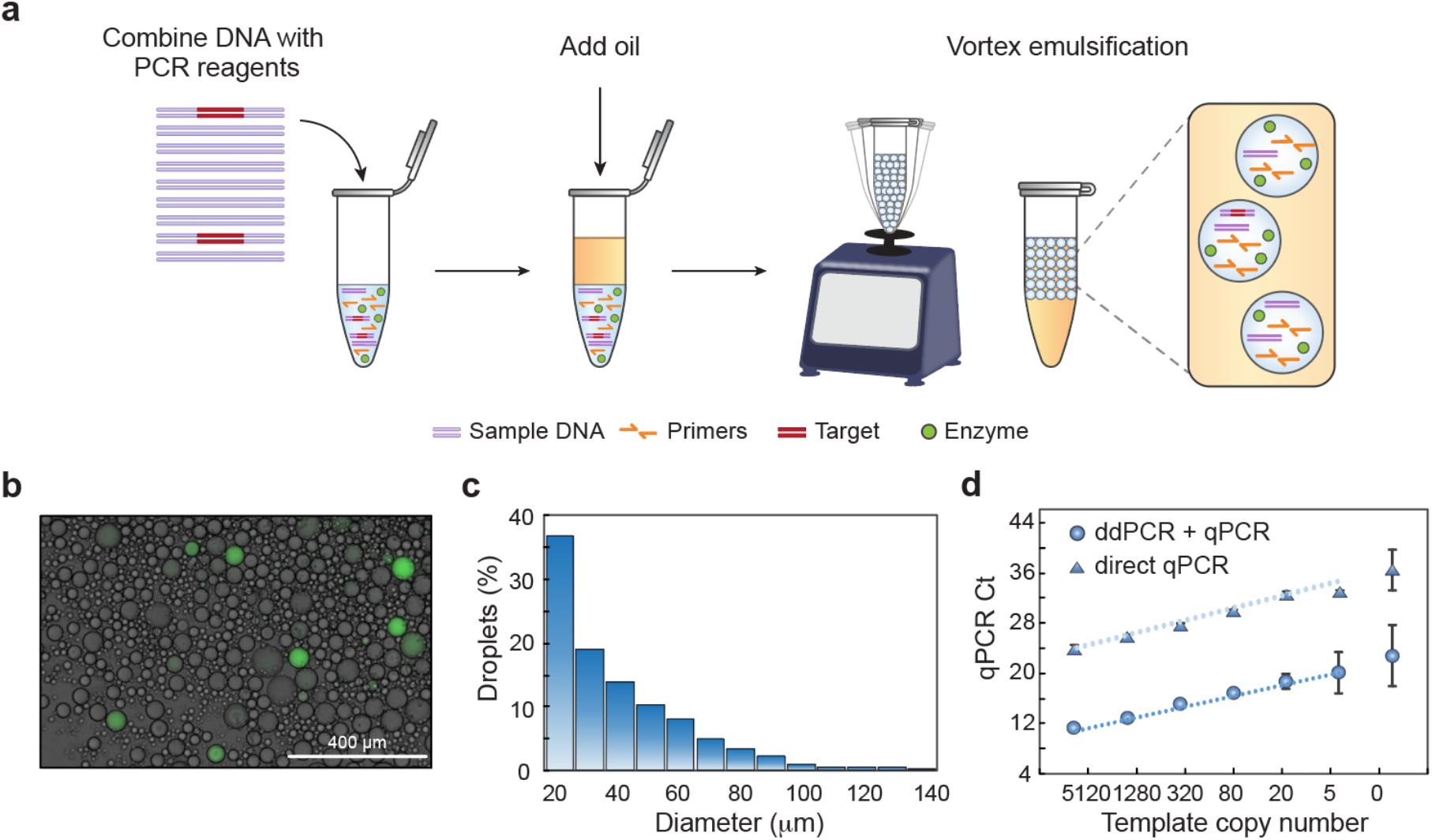
Vortex emulsification with qPCR readout enables accurate bulk ddPCR. (**a**) The DNA sample is added to a tube, oil with stabilizing surfactant is introduced, and the mixture emulsified by vortexing. (**b**) Vortexed emulsions are thermally cycled. An aliquot is amplified with TaqMan probes to enable visualization. (**c**) Size distribution of the vortex emulsified droplets obtained by imaging (*n* = 1323). (**d**) qPCR readout of vortex-emulsified ddPCR allows accurate quantitation of targets over a range of concentrations.

## Conclusions

Our approach combines the benefits of ddPCR with the simplicity, accessibility, and scalability of plate processing. We use bulk emulsification by vortexing to perform ddPCR, and bulk quantitation of generated amplicons to quantify the results, yielding a measurement accuracy superior to qPCR and similar to microfluidic ddPCR. In addition to its accuracy, our approach has benefits of ddPCR, including robustness to inhibition and efficient multiplexed reactions. Moreover, we show that a variety of amplicon detection strategies can be used with distinct advantages, such as automated electrophoresis, which is simple, fast, and accurate down to a detection limit of ∼20 molecules, and qPCR, with a sensitivity of ∼5 molecules. We also determine that, while bulk emulsified samples afford high accuracy that surpass qPCR, statistical variation in droplet size results in increased measurement error for rare targets compared to monodispersed emulsions. In instances in which this error is unacceptable, particle-templated emulsification can generate monodispersed emulsions by vortexing^14^. This approach affords other valuable features, including the ability to tune droplet size to optimize the number of amplicons generated for bulk quantitation. Moreover, using droplets of different size to analyze the same sample increases dynamic range^32^. By implementing multiplexing, it is also possible to detect a variety of targets in the same sample^28^ and to estimate the intactness of molecules based on how their subsequences co-distribute^10^, which is important for clinical diagnostics in which differentiation of fragmented and intact pathogenic genomes may be important for reducing false positive test results. In addition, multiplexing by measuring ddPCR amplicons of different length should allow simultaneous detection and quantitation of insertion, deletion, and splice mutants in research or clinical samples. Our approach thus combines key benefits of ddPCR with the simplicity and scalability of plate processing and, thus, can be readily implemented to increase the accuracy and robustness of nucleic acid testing.

## Materials and methods

### Droplet formation by commercial droplet generator

QX200 droplet generator (Bio-Rad, #1864002) was used to make emulsions following the manufacture’s instruction. Briefly, 20 μL reaction mix was prepared using ddPCR Supermix for Probe (no dUTP) (Bio-Rad, #1863024), N2 outer primers (F: AAC ACA AGC TTT CGG CAG AC, R:CCC GAA GGT GTG ACT TCC AT; final concentration of 500 nM) and template (2019-nCoV_N_Positive Control, Integrated DNA Technologies, #10006625). The ddPCR reaction mix was added to the droplet generator and converted to droplets with the use of Droplet Generation Oil for Probes (Bio-Rad, #1863005) and DG8 Cartridges and Gaskets (Bio-Rad, # 1864007).

### ddPCR and bulk readouts

Emulsified samples were transferred to PCR tubes and thermocycled in a Thermal Cycler (Bio-Rad, T100 model). Thermal cycling was performed at: 10 min at 95 ºC; 45 cycles of 20 s at 95 ºC, 30 s at 55 ºC and 30 s at 72 ºC. After ddPCR, the droplets were transferred to a flat-bottom well plate and the bulk fluorescence was measured by a Microplate reader (Tecan, Infinite 200 PRO). For BioA and Qubit measurement, 1 μL Proteinase K (800 units/ml, NEB, # P8107S) was diluted in 20 μL H_2_O and added to the thermocycled emulsions. The emulsions were then broken using 10 μL of 10% (v/v) solution of perfluoro-octanol (Sigma-Aldrich, catalog no. 370533), followed by gentle vortexing for 5 s and centrifugation for 1 min (Benchmark Scientific, MyFuge Mini centrifuge). After droplet breaking, the tubes were incubated for 10 min at 55 ºC to digest the remaining enzymes in the solution. Another incubation of 95 ºC for 10 min was used to deactivate the Proteinase K. 1 μL of the resulting solution was added directly to Bioanalyzer (Agilent 2100) or Qubit (Invitrogen, Qubit 2.0 Fluorometer) to quantify the ddPCR products. The concentration of peak of the correct molecular length was readout from Bioanalyzer. The total DNA concentration in the sample was measured by Qubit.

For qPCR readout, 1 μL of the PK treated solution was taken and diluted 100 times in DNA-free water. We used a TaqMan PCR with primers and probe targeting the ddPCR amplicon. The 20 μL qPCR reaction was assembled from 10 μL Platinum Multiplex PCR Master Mix (Life Technologies, #4464269), 1.5 μL N2 primer set (2019-nCoV RUO Kit, Integrated DNA Technologies, #10006713), 1 μL diluted ddPCR products and 7.5 μL H_2_O. The qPCR was performed in a QuantStudio 5 Real-Time PCR System (Thermo Fisher Scientific) using the following parameters: 95°C for 2 min; 40 cycles of 95°C for 15 s, 55°C for 1 min and 72°C for 30 s. Ct values for each sample was recorded as a measurement of the concentration of the target.

### Droplet formation by vortexing

DDPCR reaction mix was prepared the same as above, using ddPCR Supermix for Probe (no dUTP), N2 outprimers (final concentration of 500 nM) and template (2019-nCoV_N_Positive Control). 30 μL Droplet Generation Oil for Probes was added to the 0.2 mL PCR tube containing 20 μL ddPCR reaction mix. The tube was then placed on a vortex (Scientific Industries, digital vortex-genie 2) and agitated at 3000 rpm for 10 min. After vortexing emulsification, the samples were thermal cycled for ddPCR and readout by qPCR as described above. For positive drop visualization, one ddPCR using TaqMan primer and probe (N2 primer set) was performed. Droplets were imaged using a EVOS microscope (Thermo Fisher) with EVOS FITC LED light sources. The emulsion breaking and qPCR quantitation of ddPCR are performed the same way as above.

## Acknowledgements

Acknowledgements

We thank Joshua Batson at Chan Zuckerberg Biohub for helpful discussion. This work was supported by the Chan Zuckerberg Biohub and the National Institutes of Health (NIH) (Grant No. R01-EB019453-01 and R01-HG008978-01).

## References

1. Heid, C. A., Stevens, J., Livak, K. J., Williams, P. M., Real time quantitative PCR. Genome Res. 1996, 6 (10), 986–994.

2. Vogelstein, B., Kinzler, K. W., Digital PCR. Proc. Natl. Acad. Sci. U.S.A. 1999, 96 (16), 9236–9241.

3. Quan, P. L., Sauzade, M., Brouzes, E., dPCR: A Technology Review. Sensors 2018, 18(4), 27.

4. Heyries, K. A., Tropini, C., VanInsberghe, M., Doolin, C., Petriv, O. I., Singhal, A., Leung, K., Hughesman, C. B., Hansen, C. L., Megapixel digital PCR. Nat. Methods 2011, 8 (8), 649–U64.

5. Pohl, G., Shih, L. M., Principle and applications of digital PCR. Expert Rev. Mol. Diagn.2004, 4 (1), 41–47.

6. Alteri, C., Cento, V., Antonello, M., Colagrossi, L., Merli, M., Ughi, N., Renica, S., Matarazzo, E., Di Ruscio, F., Tartaglione, L., Colombo, J., Grimaldi, C., Carta, S., Nava, A., Costabile, V., Baiguera, C., Campisi, D., Fanti, D., Vismara, C., Fumagalli, R., Scaglione, F., Epis, O. M., Puoti, M., Perno, C. F., Detection and quantification of SARS-CoV-2 by droplet digital PCR in real-time PCR negative nasopharyngeal swabs from suspected COVID-19 patients. Plos One 2020, 15 (9).

7. Hayden, R. T., Gu, Z., Ingersoll, J., Abdul-Ali, D., Shi, L., Pounds, S., Caliendo, A. M., Comparison of Droplet Digital PCR to Real-Time PCR for Quantitative Detection of Cytomegalovirus. J. Clin. Microbiol. 2013, 51 (2), 540–546.

8. Dingle, T. C., Sedlak, R. H., Cook, L., Jerome, K. R., Tolerance of droplet-digital PCR vs real-time quantitative PCR to inhibitory substances. Clin. Chem. 2013, 59 (11), 1670–1672.

9. Hindson, C. M., Chevillet, J. R., Briggs, H. A., Gallichotte, E. N., Ruf, I. K., Hindson, B. J., Vessella, R. L., Tewari, M., Absolute quantification by droplet digital PCR versus analog real-time PCR. Nat. Methods 2013, 10 (10), 1003-+.

10. Lance, S. T., Sukovich, D. J., Stedman, K. M., Abate, A. R., Peering below the diffraction limit: robust and specific sorting of viruses with flow cytometry. Virol. J. 2016, 13.

11. Han, J., Lee, J. Y., Bae, Y. K., Application of digital PCR for assessing DNA fragmentation in cytotoxicity response. Biochim. Biophys. Acta, Gen. Subj. 2019, 1863 (8), 1235–1242.

12. Deiana, M., Mori, A., Piubelli, C., Scarso, S., Favarato, M., Pomari, E., Assessment of the direct quantitation of SARS-CoV-2 by droplet digital PCR. Sci. Rep. 2020, 10 (1), 18764.

13. Pavsic, J., Zel, J., Milavec, M., Digital PCR for direct quantification of viruses without DNA extraction. Anal. Bioanal. Chem. 2016, 408 (1), 67–75.

14. Hatori, M. N., Kim, S. C., Abate, A. R., Particle-Templated Emulsification for Microfluidics-Free Digital Biology. Anal. Chem. 2018, 90 (16), 9813–9820.

15. Baker, M., Digital PCR hits its stride. Nat. Methods 2012, 9 (6), 541–544.

16. Hatch, A. C., Fisher, J. S., Tovar, A. R., Hsieh, A. T., Lin, R., Pentoney, S. L., Yang, D. L., Lee, A. P., 1-Million droplet array with wide-field fluorescence imaging for digital PCR. Lab Chip 2011, 11 (22), 3838–3845.

17. Guo, M. T., Rotem, A., Heyman, J. A., Weitz, D. A., Droplet microfluidics for high-throughput biological assays. Lab Chip 2012, 12 (12), 2146–2155.

18. Kalinina, O., Lebedeva, I., Brown, J., Silver, J., Nanoliter scale PCR with TaqMan detection. Nucleic Acids Res. 1997, 25 (10), 1999–2004.

19. Ottesen, E. A., Hong, J. W., Quake, S. R., Leadbetter, J. R., Microfluidic digital PCR enables multigene analysis of individual environmental bacteria. Science 2006, 314 (5804), 1464–1467.

20. Shen, F., Du, W. B., Kreutz, J. E., Fok, A., Ismagilov, R. F., Digital PCR on a SlipChip. Lab Chip 2010, 10 (20), 2666–2672.

21. Suo, T., Liu, X., Feng, J., Guo, M., Hu, W., Guo, D., Ullah, H., Yang, Y., Zhang, Q., Wang, X., Sajid, M., Huang, Z., Deng, L., Chen, T., Liu, F., Xu, K., Liu, Y., Zhang, Q., Liu, Y., Xiong, Y.; Chen, G., Lan, K., Chen, Y., ddPCR: a more accurate tool for SARS-CoV-2 detection in low viral load specimens. Emerg. Microbes Infect. 2020, 9 (1), 1259–1268.

22. Morinishi, L. S., Blainey, P., Simple Bulk Readout of Digital Nucleic Acid Quantification Assays. J. Vis. Exp. 2015, (103).

23. Chantrapornchai, W., Clydesdale, F. M., McClements, D. J., Influence of relative refractive index on optical properties of emulsions. Food Res. Int. 2001, 34 (9), 827–835.

24. Liao, P. Y., Jiang, M. C., Chen, Z. T., Zhang, F. L., Sun, Y., Nie, J., Du, M. J., Wang, J. B., Fei, P., Huang, Y. Y., Three-dimensional digital PCR through light-sheet imaging of optically cleared emulsion. Proc. Natl. Acad. Sci. U.S.A. 2020, 117 (41), 25628–25633.

25. Nakayama, Y., Yamaguchi, H., Einaga, N., Esumi, M., Pitfalls of DNA Quantification Using DNA-Binding Fluorescent Dyes and Suggested Solutions. Plos One 2016, 11 (3).

26. Le Roux, D., Root, B. E., Reedy, C. R., Hickey, J. A., Scott, O. N., Bienvenue, J. M., Landers, J. P., Chassagne, L., de Mazancourt, P., DNA Analysis Using an Integrated Microchip for Multiplex PCR Amplification and Electrophoresis for Reference Samples. Anal. Chem. 2014, 86 (16), 8192–8199.

27. Bio-Rad, Droplet digital PCR applications guide. http://www.bio-rad.com/en-us/category/digital-pcr 2014.

28. Dobnik, D., Stebih, D., Blejec, A., Morisset, D., Zel, J., Multiplex quantification of four DNA targets in one reaction with Bio-Rad droplet digital PCR system for GMO detection. Sci. Rep. 2016, 6.

29. Basu, A. S., Digital Assays Part I: Partitioning Statistics and Digital PCR. Slas Technol.2017, 22 (4), 369–386.

30. Gaikwad, S. G., Pandit, A. B., Ultrasound emulsification: Effect of ultrasonic and physicochemical properties on dispersed phase volume and droplet size. Ultrason Sonochem. 2008, 15 (4), 554–563.

31. Yeung, A., Dabros, T., Masliyah, J., Czarnecki, J., Micropipette: a new technique in emulsion research. Colloids Surf., A Physicochem. Eng. Asp. 2000, 174 (1-2), 169–181.

32. Byrnes, S. A., Chang, T. C., Huynh, T., Astashkina, A., Weigl, B. H., Nichols, K. P., Simple Polydisperse Droplet Emulsion Polymerase Chain Reaction with Statistical Volumetric Correction Compared with Microfluidic Droplet Digital Polymerase Chain Reaction. Anal. Chem. 2018, 90 (15), 9374–9380.

