## Supplementary material for "Accurate bulk quantitation of droplet digital PCR": SI

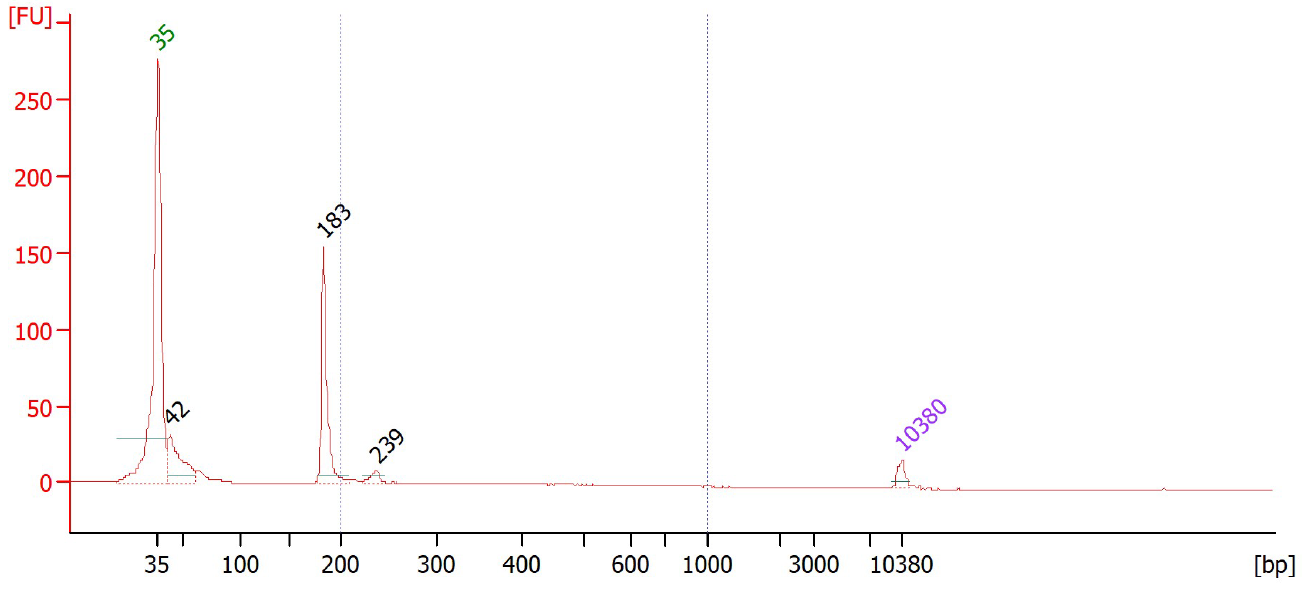

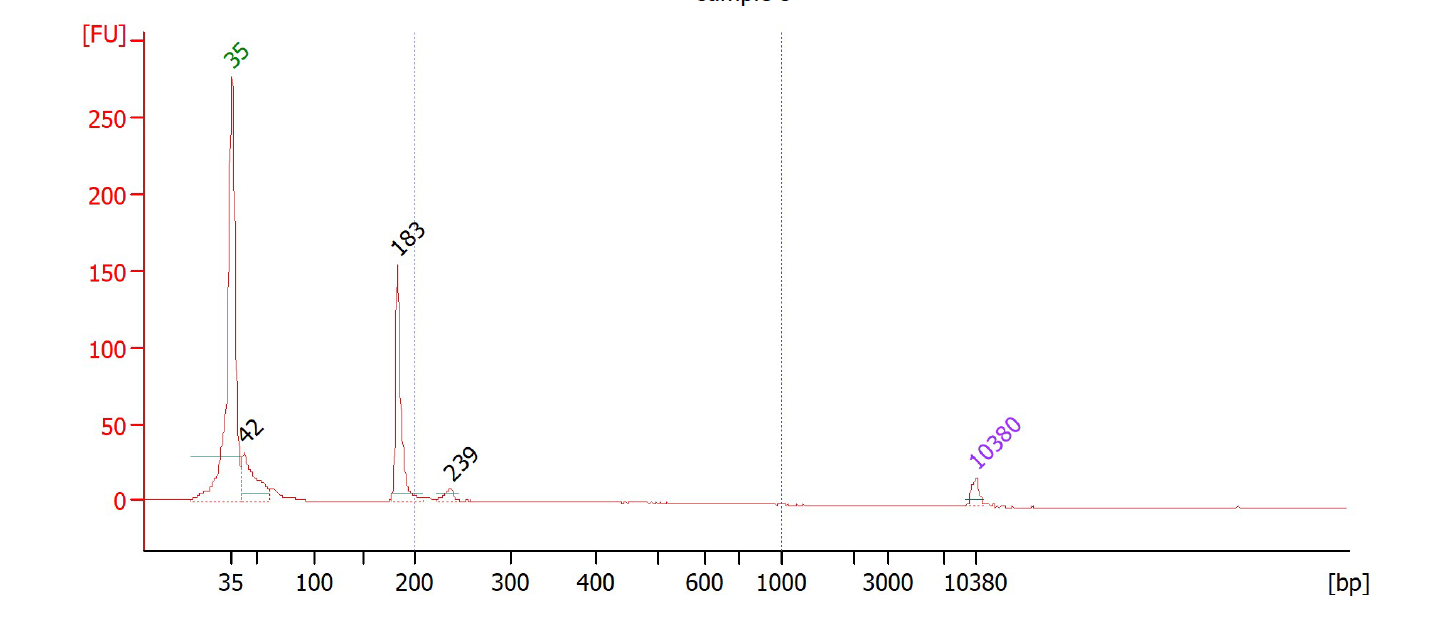

**Fig. S1** Quantification the ddPCR amplicons using on-chip electrophoresis. Peak representing the correct molecular length is detected (indicated by the arrows).
